# Spatial organization of cortical actin alignments for the ooplasmic segregation of ascidian *Ciona* eggs

**DOI:** 10.1101/849323

**Authors:** Hirokazu Ishii, Tomomi Tani

**Affiliations:** Eugene Bell Center for Regenerative Biology and Tissue Engineering, Marine Biological Laboratory, Woods Hole, MA 02543, USA; Exploratory Research Center on Life and Living Systems, National Institutes of Natural Sciences, Okazaki, Aichi 444-8787, Japan; National Institute for Physiological Sciences (NIPS), National Institutes of Natural Sciences, Okazaki, Aichi 444-8787, Japan; Biomedical Research Institute, National Institute of Advanced Industrial Science and Technology, Ikeda, Osaka 563-8577, Japan

**Keywords:** Ascidian development, Calcium wave, Cell polarity, Cytoplasmic movement, Actin, Fluorescence polarization

## Abstract

Spatial reorganization of cytoplasm in zygotic cells is critically important for establishing the body plans of many animal species. In ascidian zygotes, maternal determinants (mRNAs) are first transported to the vegetal pole a few minutes after the fertilization, and then to the future posterior side of the zygotes in later phase of the cytoplasmic reorganization, before the first cell division. Here, by using a novel fluorescence polarization microscope that reports the position and the orientation of fluorescently labeled proteins in living cells, we mapped the local alignments and the time-dependent changes of cortical actin networks in *Ciona* eggs. The initial cytoplasmic reorganization started with the contraction of vegetal hemisphere ∼20s after the fertilization induced [Ca^2+^] increase. Timing of the vegetal contraction was consistent with the emergence of highly aligned actin filaments at the cell cortex of vegetal hemisphere which ran perpendicular to the animal-vegetal axis. We propose that the first ooplasmic segregation is initiated by the local contraction of laterally aligned cortical actomyosin in the vegetal hemisphere, which in turn generates the convectional flow of cytoplasm within whole eggs.

**SUMMARY STATEMENT:** Locally distinct, transient emergence of cortical F-actin alignments were observed in live ascidian *Ciona* eggs during the first ooplasmic segregation by using fluorescence polarization microscopy.

## INTRODUCTION

During early development of many animal species including vertebrates, redistribution of cytoplasmic maternal substances in zygotes play important roles in the determination of body axes in the embryos (Lyczak et al., 2002; Ma et al., 2016; Nishikata et al., 2018; Pelegri, 2003; Weaver, 2004). The ascidian zygotes show drastic reorganization of cytoplasm from the time before the first cell division (Makabe and Nishida, 2012; Sardet et al., 2007), which has been called as the ooplasmic segregation. The initial cytoplasmic reorganization movement occurs immediately after the fertilization caused by the contraction of cortical actomyosin network (Jeffery, 1984; Jeffery and Meier, 1983; Sawada, 1983; Sawada and Osanai, 1984; Sawada and Osanai, 1985). The following reorganization of cytoplasm is driven by multiple microtubule systems that include sperm asters (Chiba et al., 1999; Goto et al., 2018; Ishii et al., 2017; Sawada and Schatten, 1988). Importantly, these cytoplasmic movements are critically important for the establishment of the body axes of ascidian embryo during the development (Makabe and Nishida, 2012; Nishida, 2005). The interplay of these two cytoskeletal dynamics helps redistribution of the maternal mRNAs called *postplasmic/PEM* RNAs, in which the transcripts of these mRNAs, including the muscle determinate *macho-1,* regulate gene expressions for cell-specific differentiations, organizing the cell polarity and cell division planes in the blastomeres (Makabe and Nishida, 2012; Nishida, 2005; Prodon et al., 2007; Sardet, 2003).

Ooplasmic segregation in ascidian eggs is triggered by fertilization. The first phase is initiated immediately after the fusion of sperm to the egg, in which the myoplasm, a mitochondria-rich cytoplasmic domain containing *postplasmic/PEM* RNAs, begins to move toward the vegetal pole. The driving mechanisms that generate the movements have remained unclear. Past studies have indicated that neither egg surface components nor inner cytoplasmic structures are responsible for the first phase of ooplasmic segregation (Jeffery and Meier, 1983; Sawada, 1983; Zalokar, 1974). EM studies have suggested that the contraction of the cortical domain (Plasma Membrane Lamina; PML) drives the first phase of ooplasmic segregation (Jeffery, 1984; Jeffery and Meier, 1983). Since this domain was stained by fluorescent phalloidin (Chiba et al., 1999; Jeffery and Meier, 1983; Sawada and Osanai, 1985), the actin cytoskeleton was likely to be the major component of this domain. Pharmacological studies using F-actin depolymerizing reagent cytochalasin B suggested the involvement of the actin cytoskeleton in the movements (Chiba et al., 1999; Reverberi, 1975; Sawada and Osanai, 1985; Takatori et al., 2015; Zalokar, 1974).

Jeffery & Meier (1983; also see Jeffery, 1984) and Sawada (1983) have proposed similar models for the mechanism of ooplasmic segregation in ascidian eggs. These models are based on the contraction of actomyosin networks that spread over the egg cortex. Their models proposed that the actin-rich cell cortex is connected to the egg plasma membrane and to inner cytoplasm. The initial event of ooplasmic segregation starts with the contraction of the basket-shape actin-rich cell cortex toward the vegetal pole. The contraction of the actin-rich cortex to the vegetal pole produces the force that displaces the vegetal cytoplasm to the animal hemisphere. The updated models also support Jeffery & Meier and Sawada models (Makabe and Nishida, 2012; Prodon et al., 2007; Sardet et al., 2007), but adding more precise explanations for the reorganization patterns of maternal factors such as mitochondria, cortical endoplasmic reticulum, and *postplasmic/PEM* RNAs during the ooplasmic segregation.

The mechanism of how actomyosin contractility leads to the directional movement of maternal substances remains an unsolved problem. We have tackled this problem by using two advanced light microscopy techniques, the selective plane illumination microscopy (SPIM) (Huisken and Stainier, 2009), and the instantaneous FluoPolScope that we have developed (Mehta et al., 2016; Nordenfelt et al., 2017; Swaminathan et al., 2017). Morphology change and the cytoplasmic movement are initiated by fertilization induced Ca^2+^ wave that is propagated from sperm fusion point (Dumollard et al., 2004). SPIM allowed to observe Ca^2+^ dynamics and the following cytoplasmic movements in whole *Ciona* egg as large as 140 µm in diameter with high signal-to-noise ratio, revealing the local differences of Ca^2+^ dynamics and cytoplasmic movement of fertilized *Ciona* egg in the vegetal hemisphere and those in the animal hemispheres in live eggs.

Alignment of cortical actomyosin filaments is the key event that determines the orientation of force production in variety of cell movements such as cell division and cell migration. The observation of molecular alignment in the cortical actomyosin network is a powerful approach to explore the mechanisms of ooplasmic segregation, but this trial has been hampered by the spatial resolution of light microscopy in the context of live cell imaging. In this study, we used the instantaneous FluoPolScope, a fluorescence polarization microscope that reports alignment of molecular assemblies with single molecule sensitivity (Mehta et al., 2016). Based on correlative imaging of the cytoplasmic movement, intracellular Ca^2+^, and the spatial organization of actin filaments in fertilized *Ciona* eggs, we found distinct spatial organizations of the actin cytoskeleton and their dynamics at the vegetal hemisphere and the animal hemisphere of *Ciona* eggs during the cytoplasmic reorganization. We will update the mechanisms of the spatial organization of cortical actomyosin for the first ooplasmic segregation in ascidian eggs.

## RESULTS

### Optical dissection of cytoplasmic reorganization in *Ciona* eggs by using selective plane illumination microscopy

We examined the dynamics of cytoplasmic reorganization in fertilized *Ciona* eggs by using a new imaging tool, the selective plane illumination microscope (SPIM). As shown in Figs. 1A,B, after the fertilization induced intracellular Ca^2+^ raise, the overall morphological change of fertilized egg was observed in parallel with the translocation of the autofluorescent myoplasm toward the vegetal pole (Deno, 1987; Nishikata et al., 1987). The myoplasm, which is composed of mitochondria-rich inner domain and cortical endoplasmic reticulum (cER) domain localized with *postplasmic/PEM* RNAs, is distributed over whole vegetal hemisphere of unfertilized *Ciona* eggs when the eggs were excited with 488nm laser light sheet illumination and observed through a green emission filter (529/24nm). Immediately after fertilization, the myoplasm moved toward the vegetal pole, which resulted in the formation of dense layer of myoplasm at the vegetal pole (Sardet et al., 1989). Together with the myoplasmic contraction, we observed a series of deformation of the fertilized egg. These morphological changes were initiated by the elevation of intracellular Ca^2+^ (Roegiers et al., 1995; Speksnijder, 1990), as was shown in the upper panel of Fig. 1B. Following the elevation of [Ca^2+^], the deformation of the egg started with a transient extension of the egg along the A-V axis for less than 2 min (aspect ratio>1), followed by a long-lasting compression to the equatorial plane (aspect ratio<1) (lower panel of Fig. 1B). The onset of the egg extension was observed ∼20s after the onset of [Ca^2+^] increase. The compression of the egg started in the middle of the slow decay phase of the fertilization induced Ca^2+^ wave.

**Figure 1.**
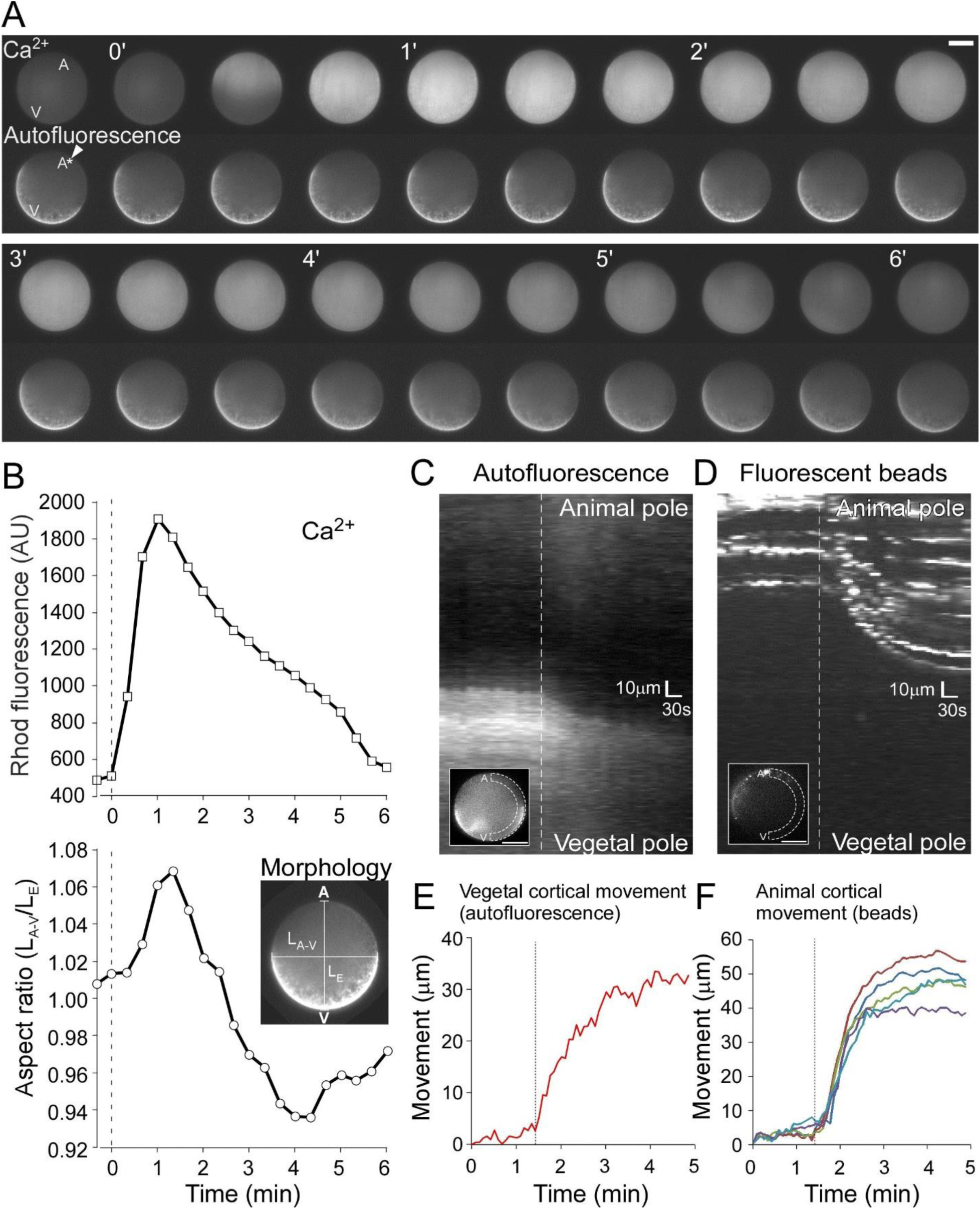
Fertilization induced intracellular Ca^2+^ wave and the following cytoplasmic reorganization in *Ciona intestinalis* egg. **A.** Time series fluorescence images of Ca^2+^ indicator Rhod-dextran (upper lane) and autofluorescence (lower lane) in fertilized *Ciona* egg. A white arrow in the first autofluorescence image indicates the position of meiotic spindle (shown as an asterisk) as a landmark of the animal pole. A and V indicate the positions of animal pole and vegetal pole, respectively. The numbers on the top left of selected images indicate the time after the start of Ca^2+^ elevation. Bar, 50 µm. **B.** Fertilization-induced Ca^2+^ changes (upper panel) and the morphology changes (lower panel) observed in **A** are plotted against time. The morphology change is expressed as a time series of change in the aspect ratio, the value derived from a length between the animal and the vegetal poles (LA-V) divided by equatorial radius (LE). Fluorescence intensities (in arbitrary unit) of Rhod-dextran are plotted against time. The time (in min) is measured after the start of Ca^2+^ elevation. **C, D.** Kymographs of autofluorescent granules (mitochondria) in the myoplasm (**C**) and 100nm fluorescent beads microinjected into the cytoplasm (**D**), both moving along the cortex of *Ciona* egg after fertilization. Averaged fluorescence intensity profiles along the curved areas enclosed by white broken lines (inset) are aligned from left to right to create the kymographs. Top of the kymograph represents the animal pole (A in the insets) and the bottom represents the vegetal pole (V in the insets). Vertical white broken lines in **C** and **D** represent the time when the movement of autofluorescent granules in the myoplasm started. **E, F.** Trajectories of autofluorescent granules in myoplasm (**E**) and 100nm fluorescent beads (**F**) that moved along the cell cortex of *Ciona* egg after fertilization. Vertical black lines in the graphs represent the time when the movement of autofluorescent myoplasm starts. The times (abscissa in min) are measured after the onset of Ca^2+^ elevation.

EM studies by Jeffery and Meier have suggested that the contraction of the egg cortical domain (Plasma Membrane Lamina; PML) drives the first phase of ooplasmic segregation (Jeffery, 1984; Jeffery and Meier, 1983). As shown in Figs. 1C,D, we monitored fertilization-triggered cortical movements in both vegetal and animal hemispheres by tracking the autofluorescent mitochondrial granules in the myoplasm and 100nm-diameter red fluorescence polystyrene beads (microsphere, ex: 580nm/em: 605nm, Molecular Probes) that had been microinjected to the cytoplasm of unfertilized *Ciona* eggs. In two kymographs along the circumference of the egg cell cortex (Figs. 1C,D), the autofluorescent mitochondrial granules at the vegetal hemisphere and the microinjected beads at the animal hemisphere both moved toward the vegetal pole along the cortex. We also observed a short, directional movement of cytoplasm with polystyrene particles in the middle of eggs, from the vegetal pole to the animal pole direction (data not shown). We found that the contraction of myoplasm at the vegetal hemisphere preceded the cortical movements at the animal hemisphere. In Fig. 1D, there was a short latent time of the start of polystyrene particle movement along the cortex at the animal hemisphere. Tracking analysis of autofluorescent myoplasm at the vegetal hemisphere (Fig. 1E) and the polystyrene particles at the animal hemisphere (Fig. 1F) revealed that the initiation of cortical movement at the animal hemisphere started approximately 5-20s after the onset of the contraction of myoplasm at the vegetal hemisphere. The total distance of translocation of microinjected beads along the cortex of animal hemisphere was approximately 40-60 µm (Fig.1F), whereas the myoplasmic movement from the equator to the vegetal pole was approximately 30 µm (Fig. 1E). The speeds at the onset of their movements were similar (58.5µm min^-1^ for myoplasm and 49.3 µm min^-1^ for beads at animal hemisphere), and gradually slowed down within 4-5 min. These SPIM observations confirmed the models by Jeffery & Meier and by Sawada, in which the initiation of myoplasmic contraction at the vegetal hemisphere triggers the first phase of ooplasmic segregation in ascidian zygotes.

### The direction of intracellular Ca**^2+^** waves correlates with the cell polarity of fertilized *Ciona* **eggs**

We observed the correlation of Ca^2+^ waves with the actomyosin-driven ooplasmic segregation and the egg morphology changes. Multiple Ca^2+^ waves with different origins occurred after fertilization. The first, fertilization induced, long-lasting large Ca^2+^ wave originated from a small area (Fig.2A, **top lane**) that had been reported as the point of sperm-egg fusion (Roegiers et al., 1995; Speksnijder, 1990). To enhance the wavefront images of the traveling Ca^2+^ waves, we processed the series of time-derivative Rhod-dextran fluorescence images by subtracting the consecutive images (taken every 2s) one after another (Fig.2A**, middle lane**). The propagating speed for the fertilization induced Ca^2+^ wave was approximately 7.5µm s^-1^, which was consistent with the previous reports (McDougall and Sardet, 1995; Speksnijder, 1990). It is worth noted that in the SPIM images in Fig. 2A, the speed of Ca^2+^ wave propagation along the cell cortex was faster than the that of the wave that ran in the middle of the cytoplasm; the shape of the wavefront was convex in the beginning, then it was straight in the middle, and then changed to concave in the end of the traveling wave.

**Figure 2.**
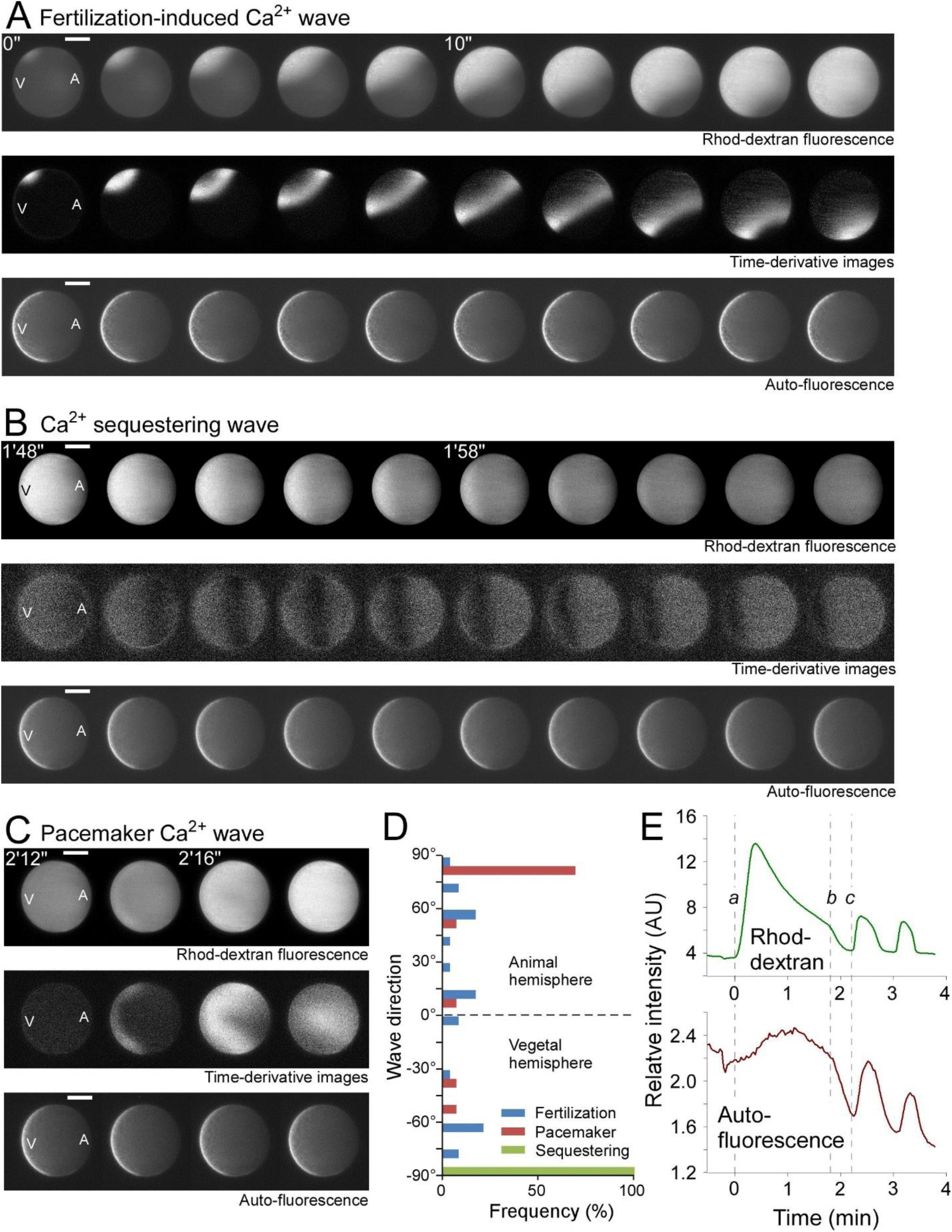
The origins and the propagation directions of intracellular Ca^2+^ waves in fertilized *Ciona intestinalis* egg. **A.** (Top lane) Time series of Rhod-dextran fluorescence images showing the propagation of fertilization induced Ca^2+^ wave. The images were taken every 200ms. Times after the beginning of initial Ca^2+^ elevation are shown at the top left of selected images. (Middle lane) Time derivative images of Rhod-dextran fluorescence obtained by subtracting two consecutive images in the top lane one after another. (Bottom lane) Time series of autofluorescence images during the first fertilization induced Ca^2+^ wave. A and V indicate the positions of animal pole and vegetal pole, respectively. Bars, 50 µm. **B.** (Top lane) Time series of Rhod-dextran fluorescence images showing the propagation of Ca^2+^ sequestering wave. Times after the beginning of initial fertilization induced Ca^2+^ elevation are shown at the top left of selected images. (Middle lane) Time derivative images of Rhod-dextran fluorescence that enhance the wavefront of Ca^2+^ sequestering wave. (Bottom lane) Autofluorescence images during a passage of Ca^2+^ sequestering wave. A and V indicate the positions of animal pole and vegetal pole, respectively. Bars, 50 µm. **C.** (Top lane) Time series of Rhod-dextran fluorescence images showing a propagation of pacemaker Ca^2+^ wave. Times after the beginning of initial fertilization induced Ca^2+^elevation are shown at the top left of selected images. (Middle lane) Time derivative images of Rhod-dextran fluorescence during the propagation of pacemaker Ca^2+^ wave. (Bottom lane) Autofluorescence images during the propagation of pacemaker Ca^2+^ wave. A and V indicate the positions of animal pole and vegetal pole, respectively. Bars, 50 µm. **D.** Histograms showing the directions of wave propagations in the first fertilization induced Ca^2+^ wave (blue), the pacemaker waves (magenta) and the sequestering waves (green). 90° represents the wave direction toward the animal pole and −90° represents the direction toward the vegetal pole. Numbers of eggs examined; 23 for fertilization induced wave, 13 for pacemaker waves and 13 for sequestering waves. **E.** Oscillation of intracellular [Ca^2+^] monitored by Rhod-dextran fluorescence (top trace) and that of autofluorescence (bottom trace). Time is measured after the beginning of initial fertilization induced Ca^2+^ elevation. Timings indicated by a, b and c correspond to the timings of the first frames of panels **A**, **B** and **C**, respectively.

A series of short repetitive Ca^2+^ waves followed the initial fertilization induced Ca^2+^ wave (Fig. 2C), which have been called as Ca^2+^ pacemaker waves (PM1, (Dumollard and Sardet, 2001; Roegiers et al., 1995; Speksnijder, 1990). In our observations, these repetitive waves originated from the vegetal hemisphere, but the initial wave was often observed not as a small point, but as a broad cup-shape area as shown in the time-derivative images (Fig. 2C, middle lane, t=2’46”). The location of this cap-shape Ca^2+^ distribution was consistent with that of autofluorescent myoplasm (Fig. 2C, bottom lane), indicating that pacemaker Ca^2+^ waves originated from the subcellular structures that associated with myoplasm. The propagation rate of the pacemaker Ca^2+^ waves was approximately 25-50 µm s^-1^, which was much faster than the fertilization induced Ca^2+^ wave.

SPIM imaging showed not only the wave of Ca^2+^ increase but also a wave of decreasing [Ca^2+^], which we called as “Ca^2+^ sequestering wave”. As far as we know, this is the first time a propagating wave of Ca^2+^ decrease is reported. The Ca^2+^ sequestering waves followed the initial fertilization Ca^2+^ wave as we clearly observed in Fig. 2B, and after the following Ca^2+^ pacemaker waves (data not shown). In other words, we observed Ca^2+^ sequestering waves every time after [Ca^2+^] increases. Interestingly, the origins of this Ca^2+^ sequestering waves were consistently from the animal pole, which is directly opposite the origin of the Ca^2+^ increase wave. This observation excluded the possibility that the Ca^2+^ sequestering wave was just a back of the Ca^2+^ wave when the wave passes by, as in many cases, we observed that the propagating direction of Ca^2+^ wave was different from that of the Ca^2+^ sequestering wave. The velocity of the Ca^2+^ sequestering waves was 5-6 µm s^-1^, which was slightly faster than the first Ca^2+^ wave and much slower than the Ca^2+^ pacemaker waves. We did not detect any difference between the velocities of Ca^2+^ sequestering waves that was observed after the fertilization induced Ca^2+^ wave and those observed after pacemaker Ca^2+^ waves. In Fig. 2D, we plotted the propagating directions of the first Ca^2+^ waves, Ca^2+^ pacemaker waves (PM1) and Ca^2+^ sequestering waves with respect to the A-V axis. The origins of the fertilization induced Ca^2+^ waves spread all over the surface of *Ciona* egg, which was not consistent with the observations using a different species of ascidian, *Phallusia*, in which the origin of the first Ca^2+^ waves was around the animal pole (Roegiers et al., 1995; Speksnijder, 1990). In the following Ca^2+^ pacemaker waves, approximately 70% (9 waves out of 13 waves) raveled to the animal pole side. In the Ca^2+^ sequestering waves, all waves we observed (13 waves) propagated along the A-V axis from the animal pole to the vegetal pole, regardless of the origins and the propagation directions of preceding [Ca^2+^] raises. Based on these observations, in later experiments, we used the directions of Ca^2+^ sequestering waves as a reference to identify the A-V axis of fertilized *Ciona* eggs.

It is worth noting that the autofluorescence of *Ciona* eggs, observed through 529/24nm emission filter, oscillated along with the Ca^2+^ waves (Fig. 2E). These changes could be observed both in the autofluorescence signal at the myoplasm as well as at the rest of the cytoplasm of the fertilized eggs. Interestingly, the changes in the autofluorescence correlated only with the Ca^2+^ pacemaker waves, but not with the first fertilization induced Ca^2+^ wave. At the onset of the fertilization induced increase in [Ca^2+^] (Fig. 2E**, vertical broken line *a***), we did not observe the increase of autofluorescence intensity, but we observed the oscillation of autofluorescence that was correlated with the Ca^2+^ oscillation.

### Correlative imaging of intracellular [Ca^2+^], cortical actin dynamics and membrane contraction of fertilized *Ciona* eggs observed with the total internal reflection fluorescence microscopy

Previous studies (Jeffery, 1984; Jeffery and Meier, 1983; Sawada and Osanai, 1984; Sawada and Osanai, 1985) have demonstrated that the first phase of ooplasmic segregation and the egg morphology changes were closely related to the contractility of actomyosin at the cell cortex of ascidian eggs. Based on fluorescence imaging of fixed cells by fluorescent phalloidin labeling (Chiba et al., 1999; Jeffery and Meier, 1983; Sawada, 1983) or by isolated cortex (Prodon et al., 2005; Sardet, 1992), the actin cytoskeleton seemed to exist very close to the cortical membrane. But the mechanisms of actomyosin system that organizes the directional movement of maternal substances toward vegetal pole was not unclear in the context of live cell imaging. We have challenged the live imaging of the actin cytoskeleton dynamics in fertilized *Ciona* eggs by using the total internal reflection fluorescence microscopy (TIRFM). We microinjected Alexa Fluor (AF) 488 phalloidin (final concentration, ∼10nM; see materials and methods for details) and Rhod-dextran (final concentration, 50-500nM; see materials and methods for details) into unfertilized *Ciona* eggs. The eggs with AF488 phalloidin and Rhod-dextran were placed on cleaned glass coverslips and were observed with TIRFM. The observed area was approximately 40-50µm in diameter, which was one third of the average diameter of *Ciona* eggs (Fig. 3A). Upon the fertilization, we observed the increases in the fluorescence intensities of both Rhod-dextran and AF488 phalloidin in *Ciona* egg cortex adjacent to the glass coverslip (Fig. 3B). Monitoring these fluorescence changes allowed us to correlate [Ca^2+^] and F-actin assembly that was labeled with AF488 phalloidin (Fig. 3C). Consistently with previous report (Dumollard et al., 2004) and with our SPIM observations (Figs. 1,2), the fertilized egg showed an initial, long lasting (∼2 min) Ca^2+^ spike, which was followed by multiple spikes with shorter durations within 5-6 min from the first [Ca^2+^] elevation. In AF488 phalloidin fluorescence signals, we observed a large transient increase followed by small fluctuations after fertilization. The onset of increase in the AF488-phalloidin fluorescence was approximately 20s-30s after the increase of [Ca^2+^] (Fig.3C; vertical lines indicate the onset times for the increases in [Ca^2+^]). These transient changes of AF488 phalloidin fluorescence always followed the transient [Ca^2+^] changes, with a short delay of approximately 20-30s for each period after [Ca^2+^] change.

**Figure 3.**
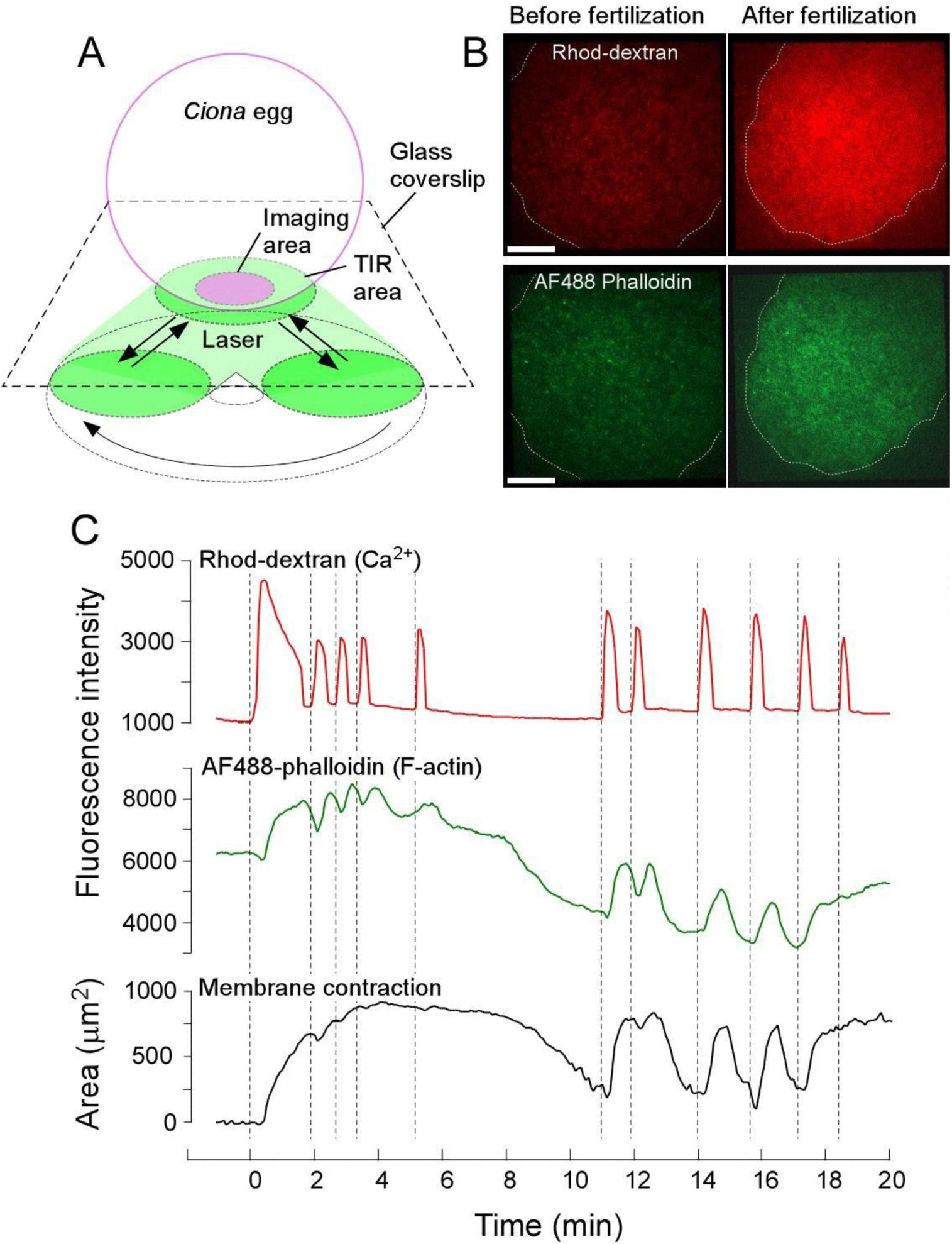
Correlative imaging of [Ca^2+^], F-actin dynamics and membrane contraction in fertilized *Ciona intestinalis* egg. **A.** A Schematic drawing showing the imaging area of *Ciona* egg placed on glass coverslip for the total internal reflection fluorescence microscopy (TIRFM). **B.** TIRF images of Rhod-dextran (Top panels) and Alexa Fluor 488 phalloidin (Bottom panels) before (left) and 30 s after (right) the onset of the first Ca^2+^ elevation. Bars, 10 µm. **C.** Time series of fluorescence intensity changes in Rhod-dextran for Ca^2+^ concentration (Top), AF488 phalloidin for F-actin (Middle) and the time-dependent reduction of membrane area attached to the glass coverslip for monitoring the membrane contraction (Bottom). Time 0 is the onset of the first Ca^2+^ elevation.

We observed the tension increase at the cortex of *Ciona* egg in the area where the egg was attached to the surface of the glass coverslip. The reduction of the attached area (Fig. 3C, lower trace, in which the y-axis value represents the *reduction of the attached area*) to the glass surface was measured correlatively with the increase of Rhod-dextran and AF488-phalloidin fluorescence, suggesting that calcium-induced actomyosin contraction induced the membrane contraction at cell surface. The onset of increase in the tension increase was approximately 20-30s after the increase of [Ca^2+^], which was consistent with the delay in the morphology change after [Ca^2+^] increase, as we observed in SPIM imaging (Fig. 1B).

Approximately 8-10 minutes after the onset of the first Ca^2+^ increase, the egg cortex entered a relaxation phase, where we observed as slacking of the membrane on the surface of the glass coverslips. AF488-phalloidin fluorescence intensity decreased to the level lower than those we had observed before the fertilization. Following this relaxation phase, we observed the second oscillation phase of [Ca^2+^] (PM2, Dumollard et al., 2004) and AF488 phalloidin labeled F-actin (staring at around 11 min in Fig. 3C) with intervals of 1-2 min. Similar to the first oscillation phase, [Ca^2+^] and AF488-phalloidin oscillations were observed in a correlated manner. For each elevation of [Ca^2+^], the increase of AF488-phalloidin fluorescence followed with a latent time of approximately 20-30s. We observed that the increase of AF488-phalloidin fluorescence and the changes in the attached membrane area occurred almost synchronously without any detectable time differences.

### Fluorescence polarization imaging of AF488 phalloidin labeled F-actin revealed the transient alignment of F-actin at the cortex of fertilized *Ciona* eggs during the first ooplasmic segregation

We used TIRFM-based instantaneous FluoPolScope (Mehta et al., 2016) for exploring the spatial/temporal dynamics of F-actin alignments at the cell cortex of *Ciona* eggs, especially during the first phase of ooplasmic segregation. In Fig.4A, we show a series of typical TIRF images of *Ciona* egg with AF488 phalloidin before (t=-1′) and after fertilization (t=3′-12′), until the time of the first cell division. In the fluorescence images (upper photographs in Fig. 4A) we observed speckles images of AF488 phalloidin bound to F-actin. We used these speckle images to compute the anisotropy of fluorescence (polarization factor) and the absolute polarization orientation of AF488 that report the degree of alignment and the net orientation of F-actin. AF488 phalloidin bound to F-actin shows the polarized fluorescence orientation that is parallel to the length of F-actin (Mehta et al., 2016). By observing the polarized fluorescence of AF488 phalloidin, we mapped the alignments of actin filaments at the cell cortex of *Ciona* eggs (lower panel of Fig. 4A). As shown in the map of fluorescence polarization orientation of AF488 phalloidin, we found highly aligned F-actin at the cell cortex at t=3′ after the fertilization. In this particular case, the orientation of F-actin alignment was around 150° in the laboratory coordinate system, as was observed in the peak of orientation histogram of the fluorescence polarization of AF488 phalloidin at t=3′ (Fig. 4B). It is worth noted that there were partially aligned F-actin to the same orientation before the fertilization, as we see the peak of the upper histogram of fluorescence polarization orientation of AF488 phalloidin in Fig. 4B. These observations suggested that there were pre-existing, weakly aligned F-actin networks at the egg cortex before the fertilization. The fertilization-induced signaling, which include the elevation of Ca^2+^, triggered the increasing density of polymerized actin (as a fluorescence intensity increase of AF488 phalloidin in Fig. 4A, upper trace) and the facilitation of F-actin alignment (as an increase of polarization factor in Fig. 4A, lower trace) to the same orientation that had existed before the fertilization. The alignment of F-actin was observed for 9-10 min after the fertilization and then disappeared along with the disassembly of F-actin (Fig. 4A, t=12′). Thus, the emergence of F-actin to the cell cortex was always associated with highly ordered alignment of F-actin as we see in the simultaneous observation of fluorescence intensity and the polarization factor of AF488 phalloidin (Fig. 4C). The polarization factor of F-actin alignment reached the highest value when the contractile ring for the first cell division at around 44-45 min after fertilization (Fig. 4A, t=44′).

**Figure 4.**
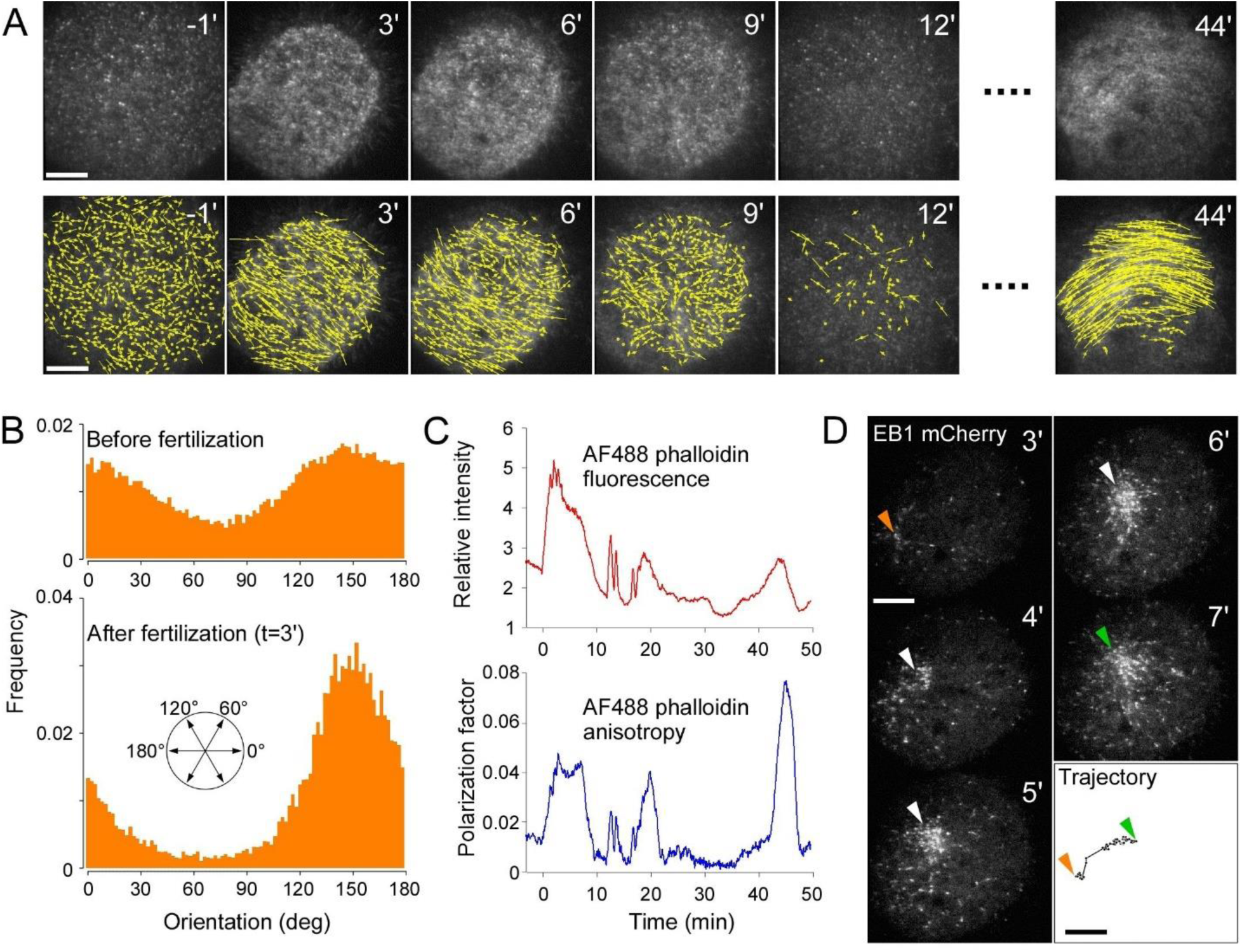
Transient F-actin alignment at the vegetal hemisphere during the first ooplasmic segregation of *Ciona intestinalis* egg. **A.** Time series of TIRF images (upper panel) and the florescence polarization orientation maps (lower panel) of a *Ciona* egg microinjected with Alexa Fluor (AF) 488 phalloidin, before and after fertilization observed at the vegetal hemisphere, as we identified from the location of sperm aster in **D**. Orientation of the yellow bars represents the fluorescence polarization orientation of AF488 and the length of the bars represents the polarization factor. Scale bars, 10µm. The numbers on the top-right indicate the time (in min) before and after the onset of initial cortical contraction of fertilized egg. **B.** Histograms of the fluorescence polarization orientation of AF488 phalloidin in *Ciona* egg cortex before the fertilization (top) and after (bottom, 3 min after the onset of the first cortical contraction) fertilization. Frequency value represents the numbers of particles divided by the total number of particles analyzed. Orientations are shown in a laboratory coordinate system. **C.** Time changes of the fluorescence intensity (top) and the fluorescence polarization of AF 488 phalloidin observed at the cell cortex of *Ciona* egg during fertilization. Time 0 represents the onset of initial cortical contraction of fertilized egg. Relative intensity is derived from the averaged fluorescence intensity of measured area divided by the averaged background fluorescence intensity of the area outside the cell. For calculating polarization factor, see materials and methods. **D.** The movement of sperm aster labeled with CiEB1-mCherry. Arrowheads indicate the locations of the aster in each image. Orange arrowhead points the initial location at t=3ʹ and the green arrowhead points the last location at t=7ʹ. Bars, 10µm.

We tried to determine the orientation of observed F-actin alignments with respect to the A-V axis. Observation of microtubules by using CiEB1-mCherry (see materials and methods for details) allowed us to correlate the movement of sperm aster with F-actin dynamics. As shown in Fig. 4D, the sperm aster, which was detected as a radially aligned CiEB1-mCherry bound to the vicinity of the plus end of microtubules, moved across the egg cortex during the cytoplasmic reorganization. Previous observations (Ishii et al., 2017; Roegiers et al., 1999; Sardet et al., 1989) have revealed that after the initial phase of ooplasmic segregation, sperm aster moves to the posterior-vegetal hemisphere along the A-V axis. The trajectory of sperm aster in Fig. 4D suggested that the alignments of F-actin that we observed in the vegetal hemisphere were normal to the A-V axis. To confirm whether the F-actin are consistently aligned perpendicular to the A-V axis in *Ciona* eggs, in the following section, we observed the alignment of F-actin at various locations where we could determine the A-V axis and the hemispheres of the observed area.

### Locally distinct cortical F-actin alignments in two hemispheres of fertilized *Ciona* eggs during the first ooplasmic segregation

Based on the correlative observation of meiotic spindle (marker of animal pole), fluorescent mitochondria (marker of vegetal hemisphere) and the Ca^2+^ wave propagations using SPIM (Figs. 1,2), we found that the A-V axis could be estimated from the orientation of Ca^2+^ sequestering wave propagation. In 14 data sets among 22 trials of the instantaneous FluoPolScope imaging of *Ciona* eggs, we correlatively observed F-actin orientations, distribution of mitochondria and yolk granules and Ca^2+^ sequestering waves before and after fertilization. We found there were two distinct F-actin alignment orientations at the cortex of fertilized *Ciona* eggs during the first phase of ooplasmic segregation. At the vegetal hemisphere where we observed a dense distribution of sub-micron diameter mitochondria (Fig. 5A) with the transmitted light imaging, the orientations of F-actin alignments were closer to the orientation normal to the A-V axis. The time series of cortical F-actin alignment changes (Fig. 5B) and the histograms of the polarization orientation of AF 488 stained F-actin before and after the fertilization (Fig. 5C) showed that in this particular case, the orientations of F-actin were around 126° (before fertilization) and 130° (after fertilization) in the laboratory coordinate system. Based on the direction of Ca^2+^ sequestering wave, the estimated A-V axis was 47°. In contrast, at the equatorial region where we observed the boundary of the region with sub-micron scale mitochondria and that with large (2-3 µm diameter) yolk granules (Fig. 5D), F-actin alignments before and after the fertilization were near parallel to the A-V axis (Figs 5G&H). We summarized the observed orientations of F-actin alignments with respect to the A-V axis at different locations before and after the fertilization of *Ciona* eggs in **Table 1**.

**Figure 5.**
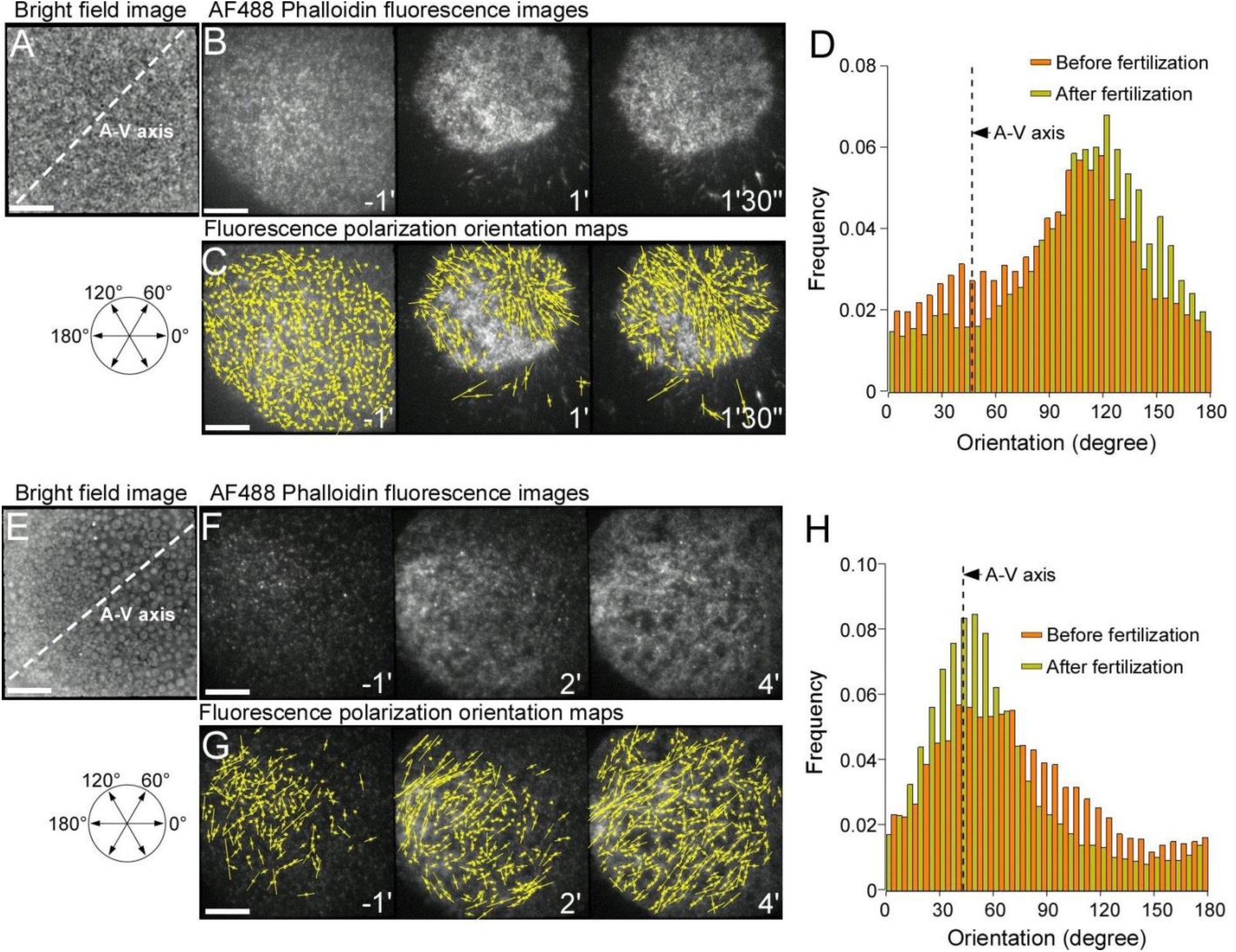
Locally distinct F-actin alignments at the animal and the vegetal hemispheres in *Ciona intestinalis* egg during the first ooplasmic segregation. **A**. Bright field image of the vegetal hemisphere of *Ciona* egg surface showing a uniform distribution of mitochondria. A-V axis was determined by the propagation orientation of Ca^2+^ sequestering waves. Bar, 10µm. **B.** Time series of TIRF images of AF488 phalloidin at the cell cortex of *Ciona* observed in the same area in **A** before (t=-1ʹ) and after (t=1ʹ, 1ʹ 30ʺ) the onset of fertilization-induced Ca^2+^ elevation. **C.** Time series of the fluorescence polarization maps of AF488 phalloidin at the cell cortex of *Ciona* in the same area observed in **A**&**B** before (t=-1ʹ) and after (t=1ʹ, 1ʹ 30 ʺ) the onset of fertilization induced Ca^2+^ elevation. **D.** Histograms of the fluorescence polarization orientation of AF488 phalloidin in *Ciona* egg before (orange) and 1min 30s after (yellowish green) the onset of fertilization induced Ca^2+^ elevation. Orientations are shown in a laboratory coordinate system. **E**. Bright field image of the lateral side of *Ciona* egg surface showing a boundary between mitochondria-rich area and that rich in yolk granules. A-V axis was determined by the propagation orientation of Ca^2+^ sequestering waves. Bar, 10µm. **F.** Time series of TIRF images of AF488 phalloidin at the cell cortex of *Ciona* in the same area observed in **E** before (t=-1ʹ) and after (t=2ʹ, 4ʹ) the onset of fertilization induced Ca^2+^ elevation. **G.** Time series of fluorescence polarization maps of AF488 phalloidin at the cell cortex of *Ciona* in the same area observed in **E**&**F** before (t=-1ʹ) and after (t=2ʹ, 4ʹ) the onset of fertilization induced Ca^2+^ elevation. **H.** Histograms of the fluorescence polarization orientation of AF488 phalloidin in *Ciona* egg before (orange) and 4min after (yellowish green) the onset of fertilization induced Ca^2+^ elevation. Orientations are shown in a laboratory coordinate system.

**Table 1.** F-actin alignments observed at various locations of cell cortex in *Ciona intestinalis* eggs. Analyses of F-actin alignments and A-V axes observed in the vegetal hemispheres (1-9), lateral sides (10-13) and in the animal hemisphere (14). Net F-actin alignments were determined from the orientation histograms of ensemble fluorescence polarization of AF488 phalloidin fluorescence. Orientation of A-V axis for each egg was identified by the propagation direction of Ca^2+^ sequestering waves observed at the same area. See materials and methods for further details.

**Ishii and Tani, Figure 6**
| Prep No. | A: Types of cortical granules | B: F-actin alignment before fertilization (degree) | C: F-actin alignment after fertilization (degree) | D: First Ca<sup>2+</sup> wave origin (degree) | E: Ca<sup>2+</sup> sequestering wave origin (degree) | F: Pacemaker Ca<sup>2+</sup> wave origin (degree) | G: Minimum angle between C and E (degree 0-90) |
| --- | --- | --- | --- | --- | --- | --- | --- |
| 1 | All mitochondria | 114 | **114** | 84 | **232** | 110 | **62** |
| 2 | All mitochondria | 9 | **12** | 224 | **81** | 254 | **69** |
| 3 | All mitochondria | 126 | **129** | 217 | **47** | 222 | **82** |
| 4 | All mitochondria | 42 | **150** | 73 | **223** | 267 | **73** |
| 5 | All mitochondria | 72 | **72** | 310 | **50** | 178 | **22** |
| 6 | All mitochondria | 78 | **78** | 33 | **73** | 40 | **5** |
| 7 | Mitochondria dominant | 48 | **52** | 323 | **131** | 227 / 67 | **79** |
| 8 | Mitochondria dominant | 108 | **117** | 65 | **56** | 255 | **61** |
| 9 | Mitochondria dominant | 93 | **96** | 143 | **152** | 323 | **56** |
| 10 | Half mitochondria and half yolk granules | 108 | **123** | 213 | **311** | 143 | **2** |
| 11 | Half mitochondria and half yolk granules | isotropic | **120** | 139 | **297** | 120 | **3** |
| 12 | Yolk granules dominant some mitochondria | 44 | **44** | 107 | **40** | 58 / 201 | **4** |
| 13 | Yolk granules dominant some mitochondria | isotropic | **81** | 289 | **220** | 38 | **41** |
| 14 | All yolk granules | 30 | **27** | 1 | **181** | 10 / 355 | **26** |

Thus, as far as based on our observations of F-actin orientation, distributions of mitochondria and yolk granules and Ca^2+^ sequestering wave, we found there was a local difference in the spatial organization of F-actin alignments with respect to the A-V axis at different locations of *Ciona* eggs during the initial phase of ooplasmic segregation. In 9 cases out of 14, at the vegetal hemisphere, the observed F-actin alignments were close to the orientation perpendicular to the A-V axis. The regions where we determined as animal hemisphere, F-actin alignments were mostly parallel along the A-V axis. We were not successful in imaging cortical F-actin orientation near animal pole and vegetal pole, presumably because the attachment of egg membrane of these areas to the glass coverslip might interfere with the normal process of ooplasmic segregation.

## DISCUSSION

### Generation of cytoplasmic movements in fertilized *Ciona* eggs during actomyosin-driven ooplasmic segregation

By using selective plane illumination microscopy (SPIM), we imaged the dynamics of cytoplasmic reorganization in *Ciona* eggs after fertilization. The primary object of this imaging was to understand the mechanisms that cause the cytoplasmic movements in fertilized *Ciona* eggs. Tracking of the movements of autofluorescent granules (mitochondria) at the vegetal hemisphere and microinjected fluorescent beads that spread over the whole cytoplasm of the eggs revealed that the onset of the first cytoplasmic reorganization was initiated by the cortical contraction at the myoplasm (Figs. 1C and E). The movement of myoplasm was followed by large cortical movements along the surface of animal hemisphere, from the animal pole to the equator along cell cortex (Figs. 1D and F). Our observations were fundamentally consistent with the hypothetical models proposed in previous studies (Jeffery, 1984; Jeffery and Meier, 1983; Sawada, 1983); the idea that the first ooplasmic segregation is based on the contraction of vegetal hemisphere and the expansion of animal hemisphere.

Simultaneous monitoring of the intracellular calcium ion concentration, [Ca^2+^], and the cytoplasmic movement revealed that in fertilized *Ciona* eggs, cytoplasmic reorganization started along with the completion of fertilization-induced [Ca^2+^] raise within the whole space of egg cytoplasm (Figs. 1A,B). In other words, the transient local difference of [Ca^2+^] during the propagation of Ca^2+^ wave was not likely to be the main cause of local difference of actomyosin-driven cytoplasmic movements during the first ooplasmic segregation. These results might be inconsistent with the previous report using *Phallusia* eggs, that there was a spatial correlation between the sperm entry site and the location of small protrusion near the vegetal pole (contraction pole) after the contraction of vegetal hemisphere (Roegiers et al., 1995; Speksnijder, 1990). Their previous observations suggested that the local difference of [Ca^2+^] during Ca^2+^ wave propagation might affect the actomyosin-driven deformation of the eggs in the vicinity of vegetal pole. Rather, our observations indicated that the spatial dynamics of the first cytoplasmic reorganization in *Ciona* eggs after fertilization is controlled by the organization of the cytoskeletal architectures that have already been programmed during the spatial organization of ooplasm during oogenesis (Prodon et al., 2006; Prodon et al., 2008).

### Roles of calcium waves for the establishment of cell polarity in fertilized *Ciona* eggs

We refined the Ca^2+^ waves in fertilized *Ciona* eggs by using the selective plane illumination microscopy (SPIM), which allowed us to analyze the propagation of Ca^2+^ waves in three-dimensional space. By tracing the wavefront of these spatial [Ca^2+^] changes, we located the origins for each wave in 3D space. Based on our analysis, we found that in *Ciona*, the origins of the first Ca^2+^ wave (fertilization Ca^2+^ wave) were not limited in the vicinity of animal pole (Fig. 2D). Series of first pacemaker waves (PM1) that follow the first fertilization induced Ca^2+^ wave, originated from a broad area within the vegetal hemisphere that corresponded to the location of the myoplasm. Observation of the vegetal origins of first pacemaker waves was generally consistent with the previous reports that the ER and mitochondria-rich domain around the contraction pole was the origins of Ca^2+^ pacemaker waves (Dumollard and Sardet, 2001; Roegiers et al., 1995; Speksnijder, 1990). However, our observations of the origins of fertilization induced Ca^2+^ waves were not consistent with the previous observations in *Phallusia* (Dumollard et al., 2003). This inconsistency might be caused by the difference of the species used.

We found a series of novel Ca^2+^ sequestering waves that followed each raise of [Ca^2+^] (Fig. 2B). Interestingly, based on our observations, the sequestering waves were consistently originated from the animal pole (Fig. 2D). The possible mechanisms for the propagating Ca^2+^ sequestration are unclear. Dumollard et al. (2003) found a Ca^2+^ sequestering function of ascidian mitochondria, which was blocked by an application of inhibitors for the mitochondrial respiration function (Dumollard et al., 2003). Similar to the simultaneous observations of [Ca^2+^] and mitochondrial flavoproteins (FAD^++^) fluorescence changes in fertilized mouse eggs (Dumollard et al., 2003), we found synchronous changes of [Ca^2+^] and autofluorescence observed through 529/24 nm emission filter (Fig. 2E), suggesting the involvement of mitochondria for the regulation of Ca^2+^ sequestration, in addition to cytoplasmic [Ca^2+^] increase. At this point, we have no evidence to suggest the intracellular signals from the animal pole that mediates Ca^2+^ sequestration waves. Since animal pole is the site for egg nucleus, some signaling molecules of nuclear origin such as cdk1-cyclinB (Bischof et al., 2017) might be involved in the Ca^2+^ sequestration.

### Local control of F-actin alignment as a possible mechanism for generating cytoplasmic streaming during the first ooplasmic segregation of fertilized *Ciona* eggs

By using our own fluorescence polarization imaging method, we have found the transient emergence and dissolution of highly aligned F-actin at the cortex of *Ciona* eggs right after the fertilization (Fig. 4A). The onset of the cortical F-actin alignments was consistent with that of the contraction of vegetal hemisphere and the expansion of cortical domains from the animal pole to the equator of the eggs, as we observed with SPIM (Fig. 1). Based on the correlative imaging of the propagation of Ca^2+^ sequestering waves that propagated along the A-V axis, we found that the orientations of F-actin alignments at the vegetal hemisphere were almost normal to the A-V axis, whereas the orientations observed at the equator and animal hemisphere were closer to the A-V axis. To our knowledge, this is the first observation of F-actin alignments and their dynamics at the cortical domain of animal eggs during fertilization. Interestingly, we observed weak alignments of cortical F-actin in unfertilized eggs, in which the orientations were similar to those observed after fertilization with respect to the A-V axis (Figs. 4&5). These observations suggest that the locally distinct spatial organizations of the actin cytoskeleton at the cell cortex of *Ciona* egg have been built during oogenesis. Thus, the basic alignments of actin filaments with respect to the A-V axis were established before the fertilization, and were kept unchanged even after the fertilization.

We found that the presence of sperm aster reorganized the alignments of F-actin as we observed less-dense and less-organized F-actin network at the location where sperm aster was observed (Figs. 5A and D). Sperm aster and associated MT system alter the morphological change of *Ciona* egg after fertilization (Roegiers et al., 1999; Sardet et al., 1989) and the following cytoplasmic rearrangement that reallocates myoplasm and associated *postplasmic/PEM* RNAs to the future posterior side of the embryo (Makabe and Nishida, 2012; Sardet et al., 2007). The roles of sperm aster for organizing the architectures of cortical F-actin have been left as unanswered important question for future studies.

Based on previously proposed models (Jeffery, 1984; Jeffery and Meier, 1983; Sawada, 1983) and our observations of F-actin alignments in *Ciona* eggs before and after fertilization, we propose a model including locally distinct F-actin alignments that might generate the force for the cytoplasmic reorganization in fertilized egg (Fig. 6). In this model we speculate two orthogonal alignments of F-actin at vegetal hemisphere and equatorial domain. At vegetal hemisphere, actin filaments align latitudinal orientation that organize the constriction of cup-shaped myoplasm along the latitudinal orientation. Like a contractile ring during cytokinesis, the contraction of laterally aligned F-actin bundles at the vegetal hemisphere squeezes the vegetal cytoplasm to the animal hemisphere. The movement of cytoplasm in the middle of egg was somehow rectified near the animal pole and was converted to the stream that moves the cortex around the animal pole toward the equator along the longitudinal orientation. This cortical flow is associated with either passive or active alignment of actin filaments along the A-V axis. We postulate that the first cytoplasmic reorganization is thus accomplished through these cortical movements associated with the locally organized contractility of actomyosin at the cortex of ascidian eggs.

**Figure 6.**
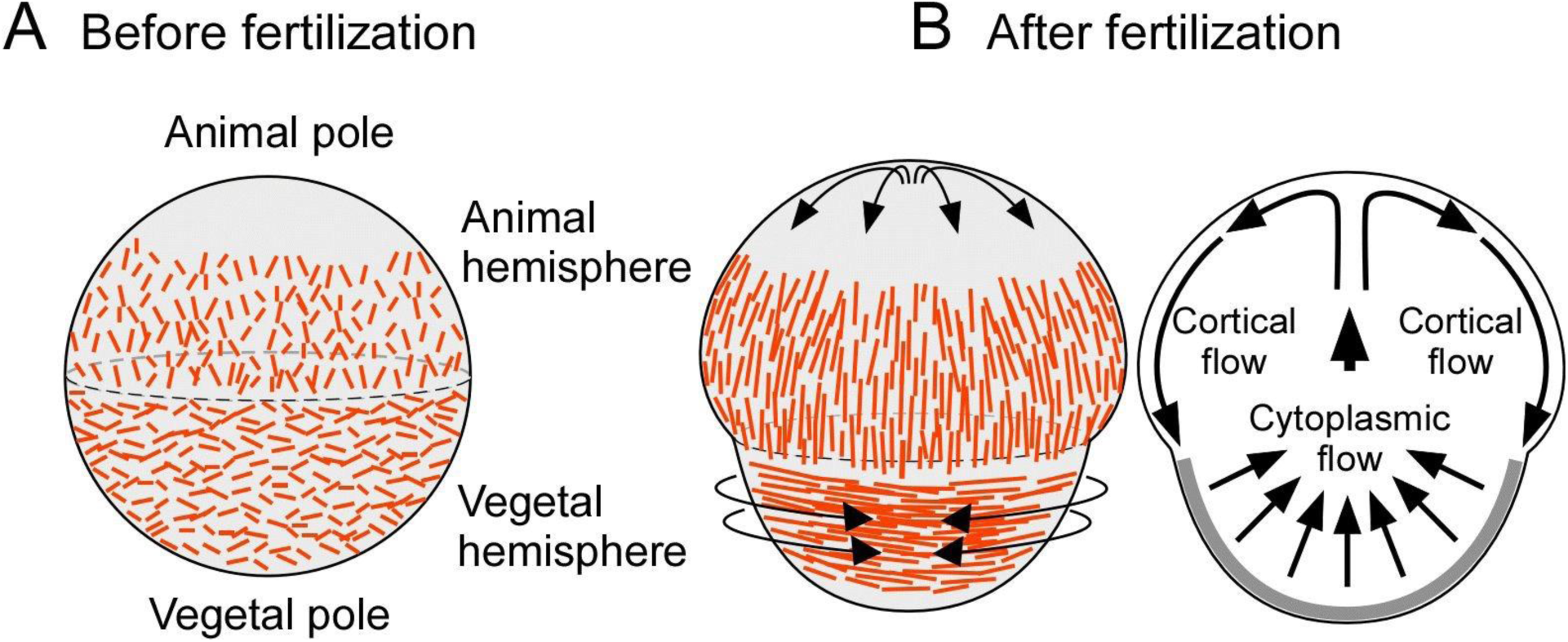
Local control of F-actin alignments for the first ooplasmic segregation. A schematic model of cortical F-actin alignment changes before (A) and after (B, left) fertilization. The contraction of actomyosin at the vegetal hemisphere and resulting compression of vegetal cytoplasm leads to the cortical flow at the animal hemisphere that reallocates materials in the cell cortex of fertilized *Ciona intestinalis* egg (B, right).

## MATERIALS AND METHODS

### Collection of animals and the preparation of eggs

The ascidia *Ciona intestinalis* were collected from local yacht harbors near the Marine Biological Laboratory, Woods Hole, Massachusetts. The eggs collected from the oviduct of dissected *Ciona* were dechorionated with 1% sodium thioglycolate (Sigma Aldrich) and 0.05% protease (from *Streptomyces griseus*, Sigma Aldrich) dissolved in filtered natural seawater (FNSW) (Ishii et al., 2014).

### Selective Plane Illumination Microscope imaging

We used a home-built selective plane illumination microscope (SPIM) for monitoring the Ca^2+^ waves and the cytoplasmic movements of *Ciona* eggs after fertilization. Laser light sheet illumination was created by scanning beams from laser heads (488nm, 20mW, Sapphire, Coherent; 561nm, 35mW; 85-YCA, Melles Griot) with electromagnetic scanning mirrors (GVS202; Thorlabs). The scanned laser beams formed a light sheet at the specimen plane through Olympus 4x 0.25NA dry objective lens. The amplitude and the scan rate of the laser beams were controlled by a dual-channel function generator (Model 4087, BK precisions). Fluorescence emission was collected through NA0.95, XLUMPLANFL 20x Olympus water immersion lens. A green filter (529/24nm; Semrock) was used for autofluorescence imaging, and a red filter (624/40nm; Semrock) was used for Rhod-dextran or fluorescent beads imaging. Fluorescence images were taken with an EMCCD camera (iXon Ultra; Andor Technology). Collected images were fed into computer (Precision 390; Dell) for further image analyses.

### Instantaneous Fluorescence Polarization imaging

The instantaneous FluoPolScope (Mehta et al., 2016) was used to observe the orientation of local F-actin. Fluorescence polarization of AF488 phalloidin speckles reports the local orientation of F-actin (Mehta et al., 2016). The laser (DPSS laser BCD; 488 nm, 20mW; Melles Griot) for excitation was circularly polarized by a quarter-wave plate (WPQ-4880–4M; Sigma-koki) and focused at the back focal plane (BFP) of the objective lens (PlanApo TIRF 60× 1.49 N.A. oil; Nikon) of an inverted microscope (TE2000-E; Nikon). The circularly polarized laser beam was rotated in the BFP of objective lens, with rotational frequency higher than 300 Hz to achieve isotropic illumination at the specimen plane. The beam was scanned along a circular annulus in the BFP using a pair of galvo-scanned mirrors (GVS202; Thorlabs). Fluorescence images were taken with an EMCCD camera (iXon+, Andor Technology). The pixel size of the EMCCD camera at the specimen plane was measured to be 180 nm. Emission filters we used were; 529/24nm transmission filter for Alexa Fluor 488 phalloidin, 609/54 nm for Rhod-dextran and for mCherry.

### Preparation of eggs for fluorescence imaging

A solution of 5 µM Alexa Fluor (AF) 488 phalloidin (life technologies) in distilled water was microinjected to the cytoplasm of dechorionated *Ciona* eggs for staining F-actin. The final concentration of AF488 phalloidin was estimated to be as low as 5-10 nM, based on the volume of microinjected AF488 phalloidin solution that was approximately from 1/1000 to 1/500 of the total cell volume of *Ciona* eggs. In general, fluorescent phalloidin as low as 10 nM has little effects on the assembly/disassembly dynamics of actin in live cells (Mehta et al., 2016; Tani et al., 2005; Yam et al., 2007). For imaging spatial/temporal changes of intracellular calcium ion concentration, we used Rhod-dextran (Molecular Probes). 50-500 µM of Rhod-dextran in distilled water was microinjected into the eggs. The final concentration of Rhod-dextran was estimated to be 50-500 nM, based on the volume of injected Rhod-dextran solution into the eggs. For imaging dynamics of microtubules in *Ciona* eggs, we constructed fluorescent protein fused microtubule plus-end binding protein EB1 from the *Ciona* EST clone (clone ID; cieg011b01) provided by Dr. Noriyuki Satoh, OIST, Japan through RIKEN BioResource Center (Satou et al., 2002). The ORF was amplified by PCR and inserted into the BmtI/BspEI site of the pRSETB-mCherry vector (Addgene plasmid 32382). The construct was transformed into the *E.coli* strain BL21(DE3). The CiEB1-mCherry proteins expressed with histidine tag were purified using a Ni-NTA column (GE Healthcare) and then the solvent was exchanged with distilled water by using NAP-5 column (GE Healthcare). Purified CiEB1-mCherry in distilled water was microinjected into the dechorionated *Ciona* eggs together with AF488 phalloidin solution. The concentration of CiEB1-mCherry was estimated to be 20-40 nM, based on the volume of microinjected CiEB1-mCherry solution and the concentration of CiEB1-mCherry (18.2 µM). For tracking the movement of cytoplasmic reorganization processes, we microinjected 100 nm diameter fluorescent polystylene beads (Molecular Probes) to the eggs. 1/100 diluted solutions (in DW) of the original 2% suspension of the beads were microinjected to the eggs. The volume of microinjection was 1/1000 to 1/500 of the total volume of *Ciona* eggs.

*Ciona* eggs with microinjected fluorescence labels/indicators were mounted on washed glass coverslips (No.1, Matsunami glass). The temperature of the imaging chamber was kept at 18°C throughout image acquisitions. For fine control of adhesion between the egg cell membrane and the glass surface, we occasionally added approximately 50 µg ml^-1^ of bovine serum albumin (Sigma Aldrich) to FNSW as needed. After letting eggs settled on the glass, we observed the eggs with transmitted light for checking the distribution of mitochondria and yolk granules. One 10^6^th dilution of sperm suspensions was applied for fertilizing dechorionated eggs.

### Image analyses

The polarization of a fluorescent particle was characterized by its degree of polarization called the polarization factor (*p*), and the orientation (*ϕ*) of its maximum polarization. The particle intensity recorded in each quadrant is related to the average intensity across the four quadrants (*I*), polarization factor, orientation (*ϕ*), and background intensity (*I*bg) as follows:

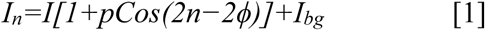

where n=0°,45°,90°,135°.

Retrieval of the polarization orientation and the polarization factor of fluorescent objects are efficiently expressed in terms of the Stokes parameters of the polarization-resolved emission:

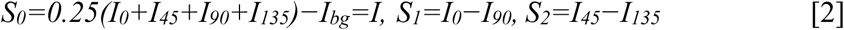

The above relationship can be written in matrix form as **S***=M(***I***−I_bg_),* where, **I**=[*I_0_, I_45_, I_90_, I_135_*]*^T^* is the column vector of input intensities, **S**=[*S_0_,S_1_,S_2_*]*^T^* is the column vector of Stokes parameters, and ***M***, called instrument matrix, is the matrix of coefficients that relate these two vectors according to Eq. [2]. The matrix ***M*** in the above equation is replaced by the instrument matrix that is generated using the tangential polarizer as described in the Appendix section of our paper (Mehta et al., 2016).

The total particle intensity, polarization factor, and ensemble orientation were obtained from Stokes parameters as follows:

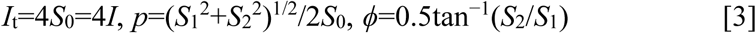

For other image analyses such as for quantitatively analyzing the propagation of Ca^2+^ waves, cytoplasmic flow and the contraction of cortical membrane areas, we used Fiji, an open source image processing package based on ImageJ (Schindelin et al., 2012). For analyzing the propagation direction of Ca^2+^ sequestering waves at the narrow egg surface observed by TIRFM, the wavefront angle was measured and the perpendicular line was defined as the wave direction. The movement of sperm aster was manually tracked by using the MTrackJ plugin (Meijering et al., 2012).

## ACKNOWLEDGEMENTS

We deeply thank Dr. Shalin Mehta for using his MATLAB codes for our fluorescence polarization analyses, and for Dr. Mark Terasaki for reading our manuscript and give us helpful feedback. We express our thanks to Dr. William Jeffery for sharing information about local *Ciona intestinalis* around the MBL. We also thank Drs. Takahito Nishikata and Takehiro G. Kusakabe for their generous supports and kind suggestions for our experiments. Our research was supported by National Institutes of Health grant R01 GM100160, Japan Society for the Promotion of Science KAKENHI grant JP18K19962 to TT, institutional funds of the Marine Biological Laboratory to TT and HI, TOYOBO Biotechnology Foundation long-term fellowship to HI.

## COMPETING INTERESTS

TT and HI have no competing interests declared.

## LEGENDS FOR SUPPLEMENTARY MOVIES

**MOVIE 01: Cytoplasmic movements in *Ciona intestinalis* egg during the first ooplasmic segregation**

Movement of microinjected fluorescent polystyrene particles (100nm in diameter, upper panel) and that of fluorescent mitochondria granules (lower panel) were simultaneously observed in fertilized *Ciona* egg with SPIM. Each frame was taken every 5s. Exposure time, 400ms. Transmittance of ND filters, 10% for 488nm 20mW laser (Sapphire, Coherent) and 5% for 561nm 35mW laser (85-YCA, Melles Griot).

**MOVIE 02: Propagation of intracellular Ca^2+^ waves in fertilized *Ciona intestinalis* egg**

Propagation of Ca^2+^ waves from the fertilization induced Ca^2+^ wave, the sequestering Ca^2+^ wave to the first pacemaker Ca^2+^ wave. Each frame was taken every 2s. Exposure time, 200ms. Transmittance of ND filter used, 5% for 561nm 35mW laser (85-YCA, Melles Griot).

**MOVIE 03: Changes of F-actin alignments at the vegetal hemisphere of *Ciona intestinalis* egg before and after fertilization revealed with fluorescence polarization microscopy**

Transient changes of F-actin alignments are shown from the time before fertilization to the first cell division of *Ciona intestinalis* egg. Positions of Alexa Fluor (AF) 488 phalloidin particles bound to F-actin are shown as yellow dots, and the fluorescence polarization orientations of the particle are shown as yellow bars. The length of the bars represents the polarization factor of the fluorescent particles. Intervals, 5s.

